# Phenotypic and Molecular Characterization of Potentially New Shigella boydii Serotype in Bangladesh

**DOI:** 10.64898/2026.11.03.763883

**Authors:** Abdus S. Mondol, Md. Pervez Hossain, Mohammad Asaduzzaman, Mohammad Aminul Islam, Mohammed Badrul Amin, Zahirul Islam, Kaisar A. Talukder

## Abstract

**Objective:** *Shigella* is a major cause of bacillary dysentery among children under 5 years of age, especially in low- and middle-income countries. At Dhaka hospital of icddr,b, every year around 3% of *Shigella* isolates from patients with diarrhea are identified as *Shigella*-like organisms (SLOs). These isolates do not react with any of the existing and provisional *Shigella* antisera. The major aims of the present study were to characterize a unique cluster of SLOs using phenotypic and genotypic methods.

**Methods:** A cluster of 6 SLOs was characterized using biochemical tests, serotyping, antimicrobial susceptibility, and virulence assay by using the Sereny test, PCR and compared their pulsed-field gel electrophoresis (PFGE) patterns with known serotypes of *Shigella* spp. Antisera against the isolate were raised in a rabbit model to determine cross-reactivity with known serotypes.

**Results:** All six SLOs were mannitol fermenting, have invasive properties, positive for *ipa*H and *Shigella* enterotoxin 2 gene along with possession of virulence plasmid of ∼ 140 MDa. All isolates were resistant to ampicillin, tetracycline, sulfamethoxazole-trimethoprim, but were sensitive to mecillinam, nalidixic acid, and ciprofloxacin. Plasmid and PFGE profiles showed that the genomic patterns of these isolates were identical but were different from all other *S. boydii* serotypes. Antiserum prepared against one of the isolates agglutinated with all isolates but did not cross-react with other *Shigella* and *E. coli* reference strains.

**Conclusion:** This unique cluster of isolates might belong to a new provisional serovar of *S. boydii* and further analysis is required to incorporate it into the stereotypic scheme of *Shigella*.

## Introduction

Shigellosis is one of the major diarrhoeal diseases in Bangladesh and several other developing countries and is responsible for a significant number of deaths especially among children less than 5 years of age [13]. *Shigella* is spread by direct fecal-hand-oral contact wherever personal hygiene is compromised [25]. Clinical infection can be transmitted by as little as 10 *Shigella* organisms [6] even without neutralization of gastric acid. The World Health Organization has emphasized the need to understand the disease burden and the epidemiology of *Shigella* infections in developing countries [4].

There are four species of *Shigella* namely, *S. dysenteriae, S. flexneri, S. boydii*, and *S. sonnei* [14] These species are further classified into 15, 23 (including subtypes), 20, and 1 serotype respectively [18]. Besides these, there are some provisional serotypes of *Shigella* and new subserotypes of *S. flexneri* [21,22,26]. This classification scheme for *Shigella* is not comprehensive, because atypical strains or newer subserotypes are being isolated in different parts of the world including Bangladesh [22]. Strains that give biochemical reactions typical of *S. boydii* but do not belong to any of the recognized O-serogroups are described as provisional serovars after rectification by a WHO International Collaborating Center for *Shigella* at CDC, Atlanta. However, their status remains provisional until several laboratories have identified such serovars and assessed their virulence properties and epidemiological importance.

The number of *S. boydii* serotypes in the *Shigella* scheme has remained at 15 since 1958. Since then, several numbers of provisional serovars of *S. boydii* have been proposed from different reference laboratories [1,9,11,26]. In 1980 [10] reported a new provisional serogroup with strain E10163, isolated in Bangladesh. Brenner and the International Committee on Systematic Bacteriology Subcommittees on the Taxonomy of *Enterobacteriaceae* (***Enterobacteriaceae*** Subcommittee of the Nomenclature Committee of the International Association of Microbiological Societies, 1958) recommended the strain be added to the scheme as *S. boydii* 18. Provisional serovar *S. boydii* E 16553 has been reported by several groups from the diverse geographical origin and designated as *S. boydii* 19 [1,11] and was also cited in the 8^th^ Edition of the Manual of Clinical Microbiology [2]. In addition, provisional *S. boydii* serovar CDC 99-4528 have been reported and designated as *S. boydii* serotype 20 [9,26].

In our previous study, we found that a significant number of strains isolated between 1999 and 2002 had biochemical properties typical of *Shigella* but could not be serotyped following the current serotyping scheme available for *Shigella* [22]. These strains were designated as *Shigella-* like *organisms* (SLO). It has been estimated that around 3% of *Shigella* infections at icddr,b hospital are caused by the SLO [21]. More importantly, the rate of isolation of these *Shigella*-like organisms in developing countries like Bangladesh has been increasing and presenting a need to completely characterize these strains. The major aims of the present study were to characterize non-agglutinating *Shigella* isolates by a battery of biochemical tests, serotyping, antimicrobial susceptibility, virulence assay by using Sereny test and test for virulence genes by PCR, and genetic fingerprinting using pulsed-field gel electrophoresis (PFGE).

## Materials and Methods

### Bacterial strains

Six isolates from 364 *Shigella*-like organisms isolated from diarrhoeal patients attending the Dhaka treatment Centre operated by icddr,b in Bangladesh were included in this study. These isolates did not agglutinate with available *Shigella* antisera and were isolated in the clinical microbiology laboratory with standard biochemical and microbiological methods [27]. All isolates were tested further in API 20E kits (bioMerieux). Strain E16553, *S. boydii* 19, was received from Central Public Health Laboratory, Colindale, London, UK. Strain CDC 99-4528 *S. boydii* 20, was received from Instiut of Pasteur, Paris, France. YSH6000, *S. flexneri* 2a [20], and an *E. coli* (ATCC 25922) strain lacking the 140 MDa invasive plasmid and sensitive to all antibiotics were used as positive and negative controls, respectively, in the Sereny test, Congo red binding ability, and in the PCR assay for detection of *ipaH* gene and *Shigella* enterotoxin genes (*set*).

### Biochemical test

Isolates were subjected to a range of biochemical tests by conventional methods as described by Ewing (1986) [8].

### Serotyping

Isolates were tested using commercially available antisera (Denka Seiken) specific for all recognized *Shigella* serovars and with antisera raised against provisional serovars *S. boydii* 19 (strains E 16553), and *S. boydii* 20 (strains CDC 99-4528). Isolates were subcultured on MacConkey agar (Difco, Becton Dickinson) plates and incubated for about 18-20 h at 37°C. Slide agglutination tests were performed as described previously [20]. Isolates were also tested in the ESYV reference laboratory with all available serogroups of *Shigella* and *E. coli*. An antiserum was raised in rabbits against a heat-killed suspension of a representative isolate (KS-49) according to the protocol of Ewing & Lindberg (1984) [7]. This antiserum was also tested against other identified SLOs, *S. dysenteriae* serotypes 13, 14, and 15, *S. boydii* serotype 19, and 20, and enteroinvasive *E. coli* serotypes O28ac, O29, O124, O136, O143, O144, and O164 for cross reactivity.

### Agglutinin-absorption tests

When cross-reactions were found, reciprocal absorptions were performed to determine the extent of the antigenic relationship between the strains [24]. Serological cross-reactions were considered significant if the heterologous titer was ≥1/16 of the homologous titer.

### Antimicrobial susceptibility

Susceptibility to antimicrobial agents was determined on Mueller-Hinton agar by the disk diffusion method following the guidelines of the Clinical and Laboratory Standards Institute [5] with commercial antimicrobial discs (Oxoid, Basingstoke, United Kingdom). The antimicrobial discs were ampicillin (10 μg), tetracycline (30 μg), mecillinum (25 μg), nalidixic acid (30 μg), sulfomethoxazole-trimethoprim (25 μg), and ciprofloxacin (5 μg). *E. coli* ATCC 25922 and *Staphylococcus aureus* ATCC 25923 were used as control strains.

### Keratoconjunctivitis assay

This test was performed according to the procedure described by Sereny (1957) [19]. Briefly, an overnight culture of bacteria, of approximately 10^10^ viable cells in 20 μl PBS, was introduced into the conjunctival sac of the guinea pig. The other eye served as the control. The guinea pigs were observed daily for 72 h and inflammatory responses were graded. The experimental procedure was approved by the animal experimental ethics committee of the icddr,b.

### Congo red binding ability

Trypticase soy broth containing 0.3% yeast extract with 1.5% agar supplemented with 0.01% Congo red (Sigma) was used to study the pigment binding ability of isolates according to the procedures of Sasakawa *et al*., (1986) [17].

### Plasmid DNA

Plasmid DNA was prepared according to the alkaline lysis method of Kado and Liu [12] with some modifications [20]. The molecular mass of the plasmid DNA bands was assessed by comparing with the mobility of the known molecular weight plasmids in *E. coli* PDK-9, R1, RP_4_, Sa, and V517 in agarose gels [20].

### Detection of virulence genes by PCR

DNA was prepared from the six isolates and examined by PCR for the presence of invasion-associated locus (*ial*) and the invasion plasmid antigen H (*ipa*H) of *Shigella* [23]. PCR to demonstrate the presence of Shiga toxin 1 (*stx*_*1*_) and *Shigella* enterotoxin (*set1* and *sen*) was performed according to published procedures [22].

### PFGE

PFGE was performed following the standardized protocol developed by PulseNet for *Shigella* [16]. Total genomic DNA was isolated in an agarose-embedded form and was subjected to enzymatic digestion with 50 U of *Xba*I. Electrophoretic separation of the DNA fragments was achieved through the contour-clamped homogenous electric field (CHEF) method on a CHEF Mapper system (Bio-Rad) using 0.5x TBE buffer (45 mM Tris-HCl, 45 mM boric acid, 1.0 mM EDTA, pH 8.0). Run conditions were generated by the auto algorithm mode of the CHEF Mapper PFGE system for sizes ranging between 30 and 600 kb, and PFGE was performed for 20 h. Analysis of the TIFF images was carried out by the BioNumerics 4.5 software package (Applied Maths, Belgium) using the Dice coefficient and UPGMA to generate dendrograms with 1.0% tolerance values.

## Results

### Biochemical characterization

All isolates showed biochemical reactions typical of *Shigella* species shown in Table 1. They were oxidase-and catalase-negative and did not ferment, arginine, lysine, sucrose, maltose, rhamnose, trehalose, raffinose, dulcitol, and sorbitol but fermented mannitol, arabinose, galactose, mannose, glucose, and xylose. API-20E tests identify all isolates as *Shigella* species.

**Table 1.** Biochemical reaction patterns of two representative strains of *Shigella*-like organisms compared with *S. boydii* 12.

| Test | Reaction |  |  |
|---|---|---|---|
|  | SLO (KS-48) | SLO (KS-49) | *S. boydii* 12 *(*K-445) |
| Iodole production | - | - | - |
| Methyl red | + | + | + |
| Voges-Proskauer | - | - | - |
| Citrate (Simmons) | - | - | - |
| H<sub>2</sub>S triple sugar iron agar | - | - | - |
| Urease | - | - | - |
| Lysine decarboxylase | - | - | - |
| Arginine dihydrolase | - | - | - |
| Ornithine decarboxylase | - | - | - |
| Motility at 37'C | - | - | - |
| Gelatin liquefaction at 22ºC | + | + | + |
| Glucose fermentation | + | + | + |
| Gas from glucose | - | - | - |
| Lactose fermentation | - | - | - |
| Sucrose fermentation | - | - | - |
| D-Mannitol fermentation | + | + | + |
| Dulcitol fermentation | - | - | - |
| Salicin fermentation | - | - | - |
| Adonitol fermentation | - | - | - |
| Inositol fermentation | - | - | - |
| Sorbitol fermentation | - | - | + |
| Arabinose fermentation | + | + | + |
| Raffinose fermentation | - | - | - |
| Rhamnose fermentation | - | - | - |
| Maltose fermentation | - | - | - |
| Xylose fermentation | (+) | (+) | (+) |
| Trehalose fermentation | - | - | - |
| Cellobiose fermentation | - | - | - |
| Galactose | + | + | + |
| Esculin hydrolysis | - | - | - |
| Melibiose | - | - | - |
| Glycerol fermentation | - | - | - |
| Acetate, sodium | - | - | - |
| Nitrate reduction | + | + | + |
| Oxidase | - | - | - |
| ONPG | - | - | - |
| MacConkey, growth in | + | + | + |
| Citrate (Christenson) | - | - | - |
| D-Mannose | + | + | + |
| Galactose | + | + | + |

+, Positive reaction within 1 or 2 days
(+), Positive reaction after 3 or more days
-, Negative reaction
ONPG, o-Nitropheny]-ß-D-galactopyranoside.

### Serotyping

All isolates were negative in agglutination tests for *Shigella* and *E. coli* serotypes and reacted only with the antiserum raised against KS-49 shown in Table 2. This was confirmed by the ESYV reference laboratory in London. Strain E16553, *S. boydii* 19, and strain CDC 99-4528 *S. boydii* 20, were also negative with the raised antiserum.

**Table 2.** Slide agglutination tests with antiserum raised against KS-49 using different serotypes of Enteroinvasive *E. coli* (EIEC) and *S. boydii* serotypes.

| Organisms tested | Slide agglutination with antiserum raised against KS-49 |
|---|---|
| Enteroinvasive *E. coli* O28ac | – |
| Enteroinvasive *E. coli* O29 | – |
| Enteroinvasive *E. coli* O124 | – |
| Enteroinvasive *E. coli* O136 | – |
| Enteroinvasive *E. coli* O143 | – |
| Enteroinvasive *E. coli* O144 | – |
| Enteroinvasive *E. coli* O164 | – |
| *S. boydii* 1 to *S.boydii* 18 | - |
| *S. boydii* provisional serotype 19 (E16553)<sup>[1]</sup> | – |
| *S. boydii* provisional serotype 20 (99-4528)<sup>[26]</sup> | – |

<sup>[1]</sup>Ansaruzzaman *et al., 2005*
<sup>[26]</sup> Woodward *et al.,* 2005

### Test for invasiveness

All isolates were positive for Congo red dye binding assay and caused keratoconjunctivitis in the guinea pig’s eye within 48 h (Table 3).

**Table 3.**
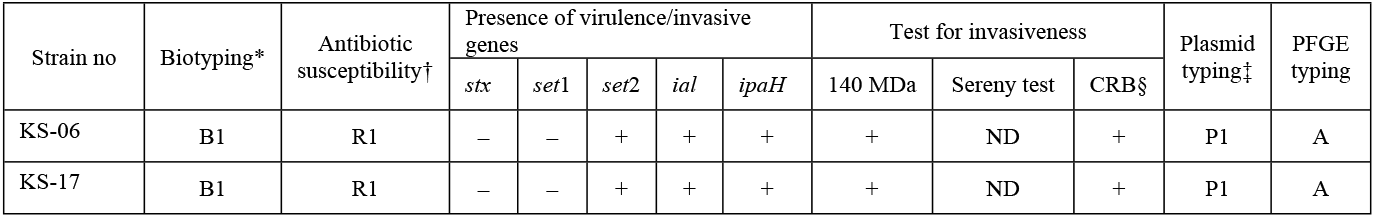
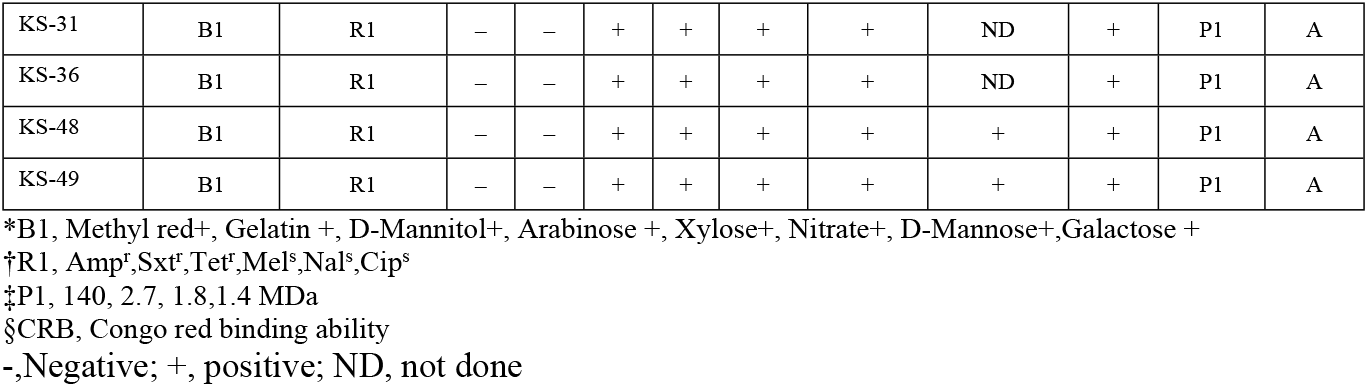
Characteristics of *Shigella-*like organisms studied.

| Strain no | Biotyping* | Antibiotic susceptibility† | Presence of virulence/invasive genes |  |  |  |  | Test for invasiveness |  |  | Plasmid typing‡ | PFGE typing |
|---|---|---|---|---|---|---|---|---|---|---|---|---|
|  |  |  | *stx* | *set*1 | *set*2 | *ial* | *ipaH* | 140 MDa | Sereny test | CRB§ |  |  |
| KS-06 | B1 | R1 | – | – | + | + | + | + | ND | + | P1 | A |
| KS-17 | B1 | R1 | – | – | + | + | + | + | ND | + | P1 | A |
| KS-31 | B1 | R1 | – | – | + | + | + | + | ND | + | P1 | A |
| KS-36 | B1 | R1 | – | + | + | + | + | + | ND | + | P1 | A |
| KS-48 | B1 | R1 | – | – | + | + | + | + | + | + | P1 | A |
| KS-49 | B1 | R1 | – | – | + | + | + | + | + | + | P1 | A |

\*B1, Methyl red+, Gelatin +, D-Mannitol+, Arabinose +, Xylose+, Nitrate+, D-Mannose+,Galactose +
†R1, Amp<sup>r</sup>,Sxt<sup>r</sup>,Tet<sup>r</sup>,Mel<sup>s</sup>,Nal<sup>s</sup>,Cip<sup>s</sup>
‡P1, 140, 2.7, 1.8,1.4 MDa
§CRB, Congo red binding ability
-,Negative; +, positive; ND, not done

### PCR assays for virulence genes

All strains were positive for *ial, ipa*H, and *sen* genes and were negative for *stx* and *set1* genes (Table 3).

### Plasmid DNA

All isolates harbored 140 MDa invasive plasmids and three other plasmids approximately 2.7, 1.8, and 1.4 MDa in size (Table 3). This plasmid pattern was different from that of other *Shigella* serotypes tested, including the provisional strains of *S. boydii* (Table 3).

### PFGE

All 6 isolates showed identical PFGE banding pattern and were closely related to the patterns of *S. boydii* 12 and were completely different from other *S. boydii* serovars (Fig 1). In addition, PFGE patterns of the tested isolates were completely different from PFGE patterns of other *Shigella* serotypes.

**Fig 1.**
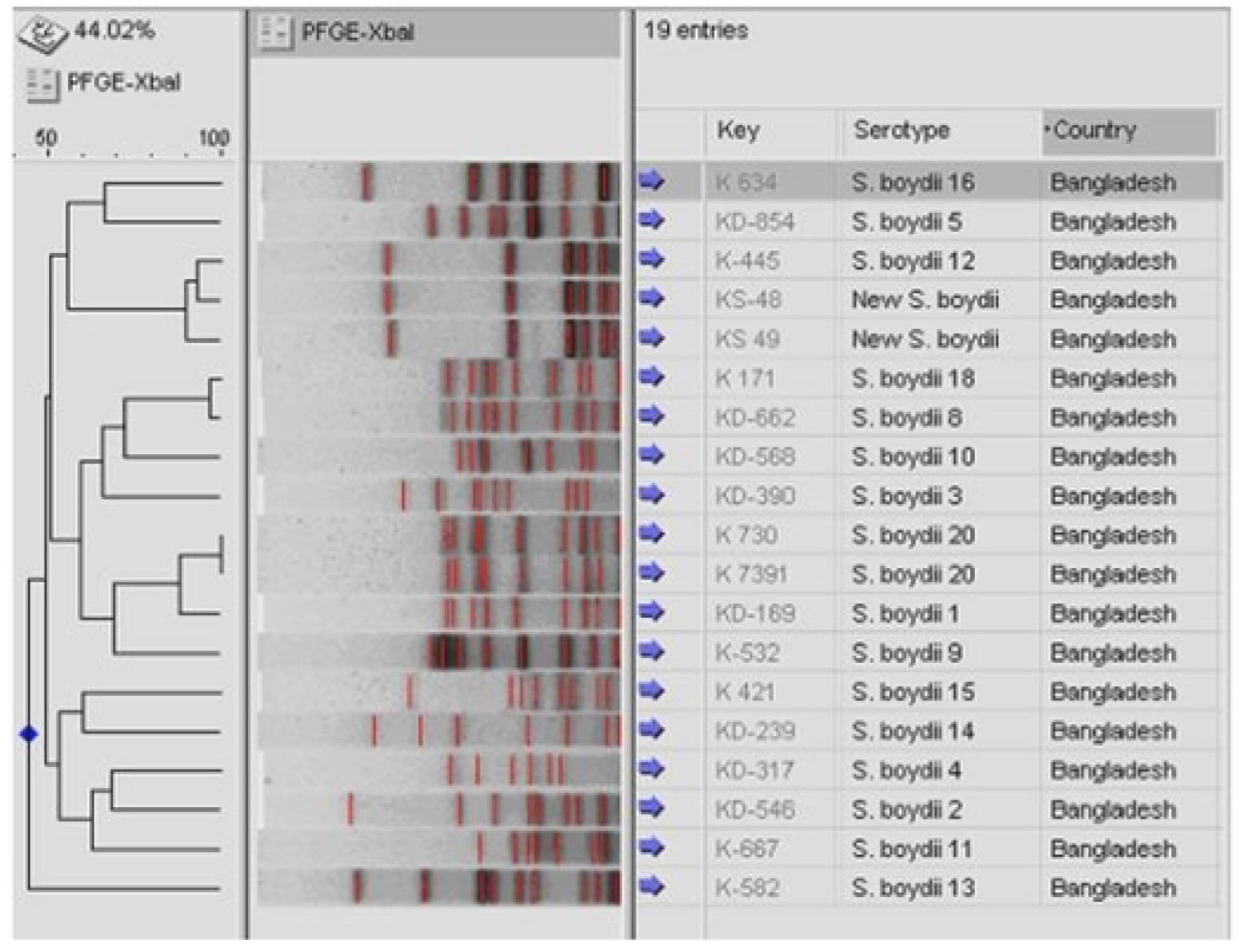
Dendrogram showing the PFGE patterns of *Shigella*-like organisms along with other existing serotypes of *S. boydii*. The dendrogram was constructed using Bionumerics 4.5 software following the unweighted pair group method with arithmetic means (UPGMA). Thin lines were added by the program to show the location (peak of densitometry curve) of less-intense bands included in the analysis.

## Discussion

The close relatedness between *E. coli* and *Shigella* species makes serological identification a crucial step in the diagnosis of *Shigella* infection. Brenner [3] considers *Shigella* organisms and *E. coli* to be a single species, based on DNA homology, and identification of a new *Shigella* serovar is very difficult due to their close relatedness. Only serological identification is not enough to identify new *Shigella* serovar. In this study, we characterized six isolates that were recovered over 3 years from hospitalized patients with diarrhea in Bangladesh (Table 3). These isolates have biochemical properties compatible with *S. boydii*.

The non-specific biochemical profile of *Shigella* serotypes and lack of criteria for discriminating between *Shigella* serotypes stresses the importance of serological identification of new *Shigella* serotypes. Therefore, antisera raised against the *Shigella*-like organisms are the best discriminating tool for differentiating *Shigella*-like organisms from the existing *Shigella* serotypes. The 6 isolates were found to react only with the antisera raised against KS-49. Reciprocal agglutinin-absorption tests showed that all the isolates were identical. These appear to be new provisional serotypes of *S. boydii* belonging to a yet uncharacterized antigenic determinant.

Two novel toxins, *Shigella* enterotoxin 1 (ShET-1) encoded by chromosomal gene *set*1 and *Shigella* enterotoxin 2 (ShET-2) encoded by *sen* (located in 140 MDa plasmid), have been associated with the pathogenicity of many *Shigella* species [15]. The positive results from invasiveness assays such as the Sereny test, presence of 140 MDa plasmid, *Shigella* enterotoxin (*set* 2) gene, *ial*, and *ipaH* genes provide evidence that these isolates have virulence properties that are typical of *Shigella* species. However, EIEC are also positive for the same virulence properties. The identity of tested strains as *Shigella* was confirmed based on the biochemical tests which predominantly matched with *S. boydii*.

Although none of six strains was positive for agglutination assays with any serovar of *S. boydii*, due to the matching biochemical reaction patterns with *S. boydii* we assumed that these strains might be new variants of *S. boydii*. To exclude the possibility that these isolates are genetically linked with existing serotypes of *S. boydii* we compared the PFGE banding patterns of these isolates with other serotypes of *S. boydii*. The results had clearly shown that all the tested isolates had PFGE patterns closely related to *S. boydii* 12 but completely different from that of any of the other *Shigella boydii* serotypes (Fig 1). PFGE does not have the sensitivity to precisely identify genetic differences between two closely related strains, hence further analysis of the isolates by using whole genome sequencing would be helpful to get more insights into the level of differences between these isolates. Further, antigenic variations between these isolates and the existing serotypes need to be analyzed at the genomic level to confirm the serotype differences. Nevertheless, this study provides valuable data on the identification and characterization of a potentially new serovar of *Shigella* spp which could be the foundation of future studies leading to confirmation and nomenclature of this potentially new serovar.

## Conclusion

In summary, the SLO isolates characterized in this study exhibited biochemical, serological, and molecular traits related to the genus *Shigella*. No serological cross-reactivity was observed with other *Shigella* and *E. coli* serotypes. These isolates are likely to possess an antigenic epitope that makes this group of isolates different from all other recognized serotypes of *Shigella* and therefore we propose that these isolates represent a novel emerging serovar of *S. boydii*.

## Authors’ Declaration Statements

### Ethics approval and consent to participate

Not applicable

### Availability of data and material

The data used in this study are available and will be provided by the corresponding author upon a reasonable request.

### Authors’ Contributions

The research team designed and led the design of the study and the preparation of the article; also carried out data extraction, analysis, and review of the literature for data retrieval. The final version of this manuscript was read and approved by all authors. All of the authors got input and helped shape the research, review, and manuscript.

## Acknowledgment

This research was funded by the Government of the Peoples Republic of Bangladesh. icddr,b acknowledges with gratitude the commitment of the Natural Environment Research Council (NERC), the UK to its research efforts. icddr,b is also grateful to Canada, Sweden, and the UK for providing core/unrestricted support.

